# The influence of age and sex on the pre-treatment immune microenvironment of a carcinogen induced murine model of bladder cancer

**DOI:** 10.1101/2021.09.10.459788

**Authors:** Ali Hamade, Deyang Li, Kathrin Tyryshkin, Minqi Xu, Stephen Chenard, Gwenaelle Conseil, Priyanka Yolmo, D. Robert Siemens, Madhuri Koti

**Affiliations:** Queen’s Cancer Research Institute, Kingston, ON, Canada; Department of Biomedical and Molecular Sciences, Queen’s University, Kingston, ON, Canada; Department of Pathology and Molecular Medicine, Queen’s University, Kingston, ON, Canada; Department of Urology, Queen’s University, Kingston, ON, Canada

**Keywords:** Bladder cancer, Tumor immune microenvironment, B cells, Tertiary lymphoid structure, BBN carcinogen, Murine model

## Abstract

The incidence of urothelial carcinoma of the bladder is four times higher in males than females; however, females tend to present with a more aggressive disease, a poorer response to immunotherapy and suffer worse clinical outcomes. Recent findings have demonstrated sexual dimorphism in the tumor immune microenvironment of non-muscle invasive bladder cancer and associated clinical outcomes. However, a significant gap in knowledge remains with respect to the current pre-clinical modeling approaches and more precisely recapitulating these differences towards improved therapeutic design. Based on the similarities in mucosal immune physiology between humans and mice, we evaluated the sex and age-related immune alterations in healthy murine bladders. Bulk-RNA sequencing and multiplex immunofluorescence based spatial immune profiling of healthy murine bladders from male and female mice of age groups spanning young to old showed a highly altered immune landscape that exhibited sex and age associated differences, particularly in the context of B cell associated responses. Spatial profiling using markers specific to macrophages, T lymphocytes, B lymphocytes, activated dendritic cells, high endothelial venules, myeloid cells and the PD-L1 immune checkpoint showed sex and age associated differences. Bladders from healthy older female mice showed a higher number of tertiary lymphoid structures (TLSs) compared to both young female and male equivalents. Spatial immune profiling of N-butyl-N-(4-hydroxybutyl) nitrosamine (BBN) carcinogen exposed male and female bladders from young and old mice revealed a similar frequency of TLS formation, sex differences in the bladder immune microenvironment and, age differences in latency of tumor induction. These findings support the incorporation of sex and age as factors in pre-clinical modeling of NMIBC and will potentially advance the field of immunotherapeutic drug development to improve clinical outcomes in NMIBC.

## Introduction

Bladder cancer accounts for 200,000 annual deaths globally^1^ and presents as non-muscle invasive bladder cancer (NMIBC) or muscle invasive bladder cancer (MIBC). Bladder cancer remains the most management intensive and expensive cancer to treat in the elderly. Because of the high tumor mutational burden, bladder tumors exhibit high immunogenicity. Patients with intermediate or high risk NMIBC receive intravesical therapy with an attenuated strain of Mycobacterium bovis (*M.bovis*) also known as bacillus Calmette-Guérin (BCG; commonly used as a vaccine for prevention of tuberculosis).^2^ Despite the use of BCG immunotherapy as a gold standard for over 45 years, up to 50% patients exhibit poor responses, with many suffering from recurrence and progression to MIBC. Recent immune checkpoint blockade therapies have shown modest benefits in patients with bladder cancer.

With an average age of 73 years at diagnosis, bladder cancer exhibits four times higher incidence in males compared to females. Female patients with NMIBC generally present late and display earlier recurrence and more advanced tumor stages than males.^3–6^ Bladder cancer also exhibits significant sex disparity in outcomes following immunomodulatory treatments.^3,7,8^ Several previous reports have shown the influence of hormones and sex-chromosome associated genetic links mediating such sexual dimorphism; however, the influence of immunological factors remain less studied. With biological aging, the immune system experiences significant decline in function. Interestingly, this phenomenon, also known as immunosenescence, progresses in a sex-differential manner.^3,9–12^ The bladder is an organ with a mucosal barrier providing protection from uropathogens and therefore exhibits unique immunological characteristics that sustain a local physiological microbiome.^13–15^ Simultaneous to changes in peripheral immune profiles, significant alterations within the mucosae also occur with biological aging and were recently reported in the context of healthy female murine bladder.^16^

While the significance of age and sex as factors associated with incidence and clinical outcomes in bladder cancer have been well acknowledged,^3,7,8^ current pre-clinical modeling approaches do not incorporate these variables in experimental design. Furthermore, although the widely used orthotopic cancer cell implantation based syngeneic models are useful in addressing the mechanistic interactions associated with carcinogenesis and/or response to targeted therapies, these may not fully reflect the events associated with local and systemic immune physiological shifts and functional alterations that accompany biological aging and those that are sex-dependent. This is more critical in the context of BCG immunotherapy in the elderly, where it also acts as an adjuvant for both innate and adaptive immune compartments. The variabilities in cell numbers used at implantation may also affect the observed response in murine models. Importantly, due to the ease and consistency of catheterization procedures, female mice are generally used in most of the bladder cancer implantation models. Exposure to the carcinogen N-butyl-N-(4-hydroxybutyl) nitrosamine (BBN), a mutagen mimicking cigarette smoke, recapitulates the most common risk factor in bladder cancer.^17^ This widely used bladder cancer model also exhibits tumor mutational spectrum and heterogeneity and has been employed for investigations in both NMIBC and MIBC.^18, 19^ Several additional genetically engineered and transgenic models of bladder cancer have been developed; however, their age and sex associated local immune physiological alterations are yet to be fully characterized.

In the current study we first characterized the age and sex associated bladder immune landscape alterations in physiologically healthy female and male mice. We then characterized the sex and age associated pre-treatment bladder immune microenvironment in the BBN induced syngeneic murine model of bladder cancer.

## Materials and methods

All murine experimental procedures were approved by the Queen’s University Animal Care Committee. Physiologically healthy C57BL6/J male and female mice belonging to age groups ranging from young to old were obtained from Charles River Laboratory (Montreal, QC, Canada). Mice were fed a sterilized conventional diet and water and maintained under a pathogen-free environment.

### Bulk-RNA sequencing and gene set enrichment analysis

Whole bladders were collected from 3-, 6-, 9-, 12-, 15- and 18-month-old healthy female and male mice (n=3-4 each group and sex) and snap frozen in liquid nitrogen. These age groups were selected to mimic the young (3-6 months), middle (9-12) and older (15-18) ages in mice, as previously defined.^20^ Total RNA from fresh-frozen bladders was isolated using the RNeasy Mini Kit (Qiagen Inc, Canada). Qualitative and quantitative assessments of RNA were performed using a spectrophotometer (Nanodrop ND-100) and a bioanalyzer (Agilent). Samples that met the purity requirement (>8.0 RNA integrity number) were subjected to bulk RNA-sequencing (Genome Quebec, Montreal, QC, Canada). RNA-sequencing data analysis was performed using previously established protocols.^21^ Pre-ranked lists resulting from DeSeq2 analysis were subjected to Gene Set Enrichment Analysis (GSEA; http://software.broadinstitute.org/gsea/) for comparison against GO Biological Processes gene sets for a preliminary gene ontology analysis (version 7.4). Findings were validated using a proprietary feature selection algorithm (Tyryshkin K., *Unpublished*) after TPM normalized counts were filtered to isolate genes with the top 10% of expression to find the top 1% of differentially expressed genes between comparison groups. Top 200 differentially expressed genes (identified via DeSeq2 analysis) from each comparison within and between sexes were also subjected to further analysis to identify enrichment of genes predicting cell subsets using the MyGeneset application (https://www.immgen.org/).^22,23^ Details of additional analysis are provided in supplementary methods.

### BBN carcinogen treatment

Female and male C57BL/6J mice belonging to young (5 months) and middle (12 months) and older (15 months) age groups (n=15-20 each age group) were administered 0.05% N-Butyl-N-(4-hydroxybutyl) nitrosamine (BBN; TCI America) *ad libitum* in drinking water (protected from light) once a week for 12 weeks as per previously established protocols.^24^ Ultrasound based imaging was performed at 4-week intervals to monitor changes in the bladder wall. Bladders were collected at 4, 7- and 12-weeks post initiation of BBN exposure to characterize the bladder immune microenvironment by histopathology and multiplex immunofluorescence (IF).

### Multiplex immunofluorescence staining

Following histological evaluation using hematoxylin and eosin (H&E) staining at the Queen’s Molecular Pathology Laboratory, paraffin embedded tissue sections (4 μm) were subjected to spatial immune profiling using multiplex IF staining to analyze infiltration patterns of selected immune markers using two panels of antibodies to identify CD11b+ myeloid cells, CD3+ total T cells, CD8+ cytotoxic T cells, PNAd+ high endothelial venules, Pax5+ B cells, PD-L1 immune check point, CD163+ M2-like macrophages, EpCAM+ epithelial cells (Supplementary Table 1). Staining was performed at the Molecular and Cellular Immunology Core (MCIC) facility, BC Cancer Agency as per previously reported methods.^8^ Multiplex IF stained images were acquired using the Vectra 3 multichannel imaging system. Images were then imported to Phenochart (1.1.0) and Inform Viewer (2.5.0) software for annotation followed by capturing at high magnification. The Inform Viewer (version 2.5.0, Akoya Biosciences) was used to visualize a detailed composite image of each section with the flexibility of spectral separation of all the fluorophores from each panel. Immune cell infiltration was evaluated in 3-5 random high-power fields adjacent to the basal layer of the urothelium in the bladders from healthy and BBN treated mice.

## Results

### Bulk-RNA sequencing reveals age and sex associated differences in transcriptome profiles of healthy murine urinary bladders

We characterized the local immune microenvironment of the healthy murine urinary bladders to determine whether any age associated differences existed between females and males. To measure the age-related transcriptomic alterations in the bladder immune microenvironment, we first performed an unsupervised feature selection following the preprocessing of the RNA-seq files. Unsupervised heatmap clustering using the top 25% ranked genes, automatically distinguished the female and male sexes in all age groups (representative comparisons from two age groups are shown in Supplementary Figure 1). These findings confirmed that sex differences in murine urinary bladder are present across different ages.

### B cell-function associated pathways are enriched in bladders from healthy aged females compared to aged males

Pathway enrichment was performed with the gene set enrichment analysis (GSEA) tool using whole transcriptomic log2fold change ranked lists obtained from DESeq2 analysis (Supplementary Table 2, 3 and 4). GSEA analysis revealed enrichment in pathways associated with B cell receptor signaling and B cell mediated immunity among the top 10 enriched pathways starting at 12 months of age (Figures 1A-1C). While similar enrichment was observed upon comparison of the 18 months old group with the 3 months old group within both sexes, between sex comparisons revealed significant increase in enrichment score in 18-month-old female compared to their male counterparts (Figure 1A). The 12-months, 15 months, and 18 months old females had a significantly greater enrichment (*q-value* <0.05, false discovery rate; FDR) of B cell and other immune-related biological processes compared to 3-month-old female mice (Figure 1B). Cytoscape analysis of all significantly upregulated GO pathways within and between sexes in the aged vs young context (*q*-*value*<0.05, FDR), further confirmed that multiple pathways associated with immune response and B-cell function cluster together with similar age associated trends within and between sexes (Supplementary Figure 2).

**Figure. 1.**
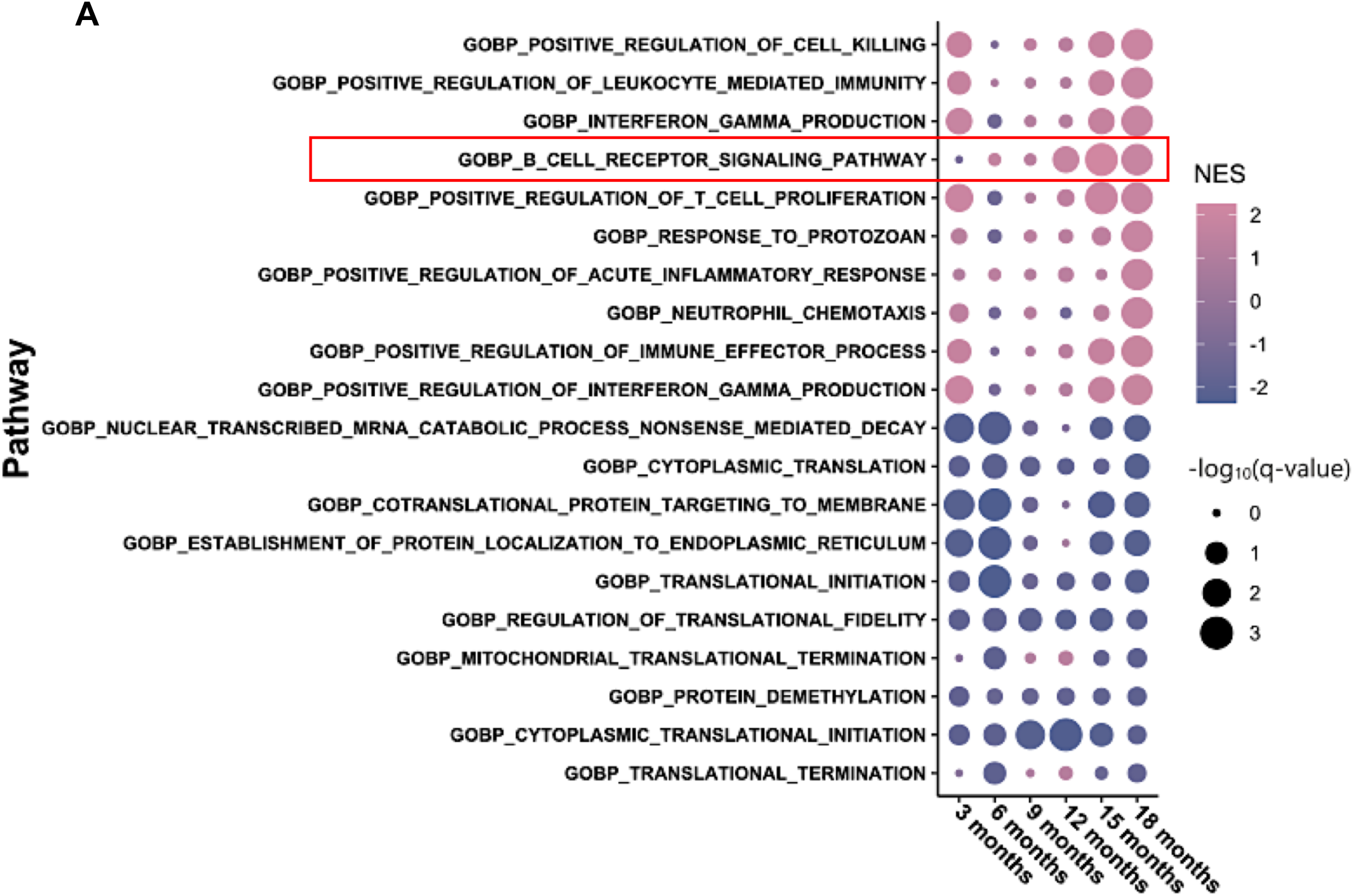
Urinary bladder of healthy C57BL/6 mice exhibit age and sex-associated enrichment in immune response pathways. Enrichment of the top 10 significantly enriched and downregulated GO biological processes pathways in **A)** females compared to age-matched males, **B)** aged females compared to 3-month-old females and, **C)** aged males compared to 3-month-old males. Total RNA from healthy male and female urinary bladders from different age groups was subjected to bulk-RNA sequencing. Differential expression was assessed using DESeq2 in R (1.4.0). GSEA pre-ranked analysis of whole transcriptomic Log2foldchange ranked lists from DESeq2 was performed. Plots were generated using ggplot2 (3.3.5), with dot size corresponding to -log10(q-value, FDR), and the color associated with normalized enrichment score (NES). Red box highlights the B cell receptor signaling pathway.

**Figure. 1.**
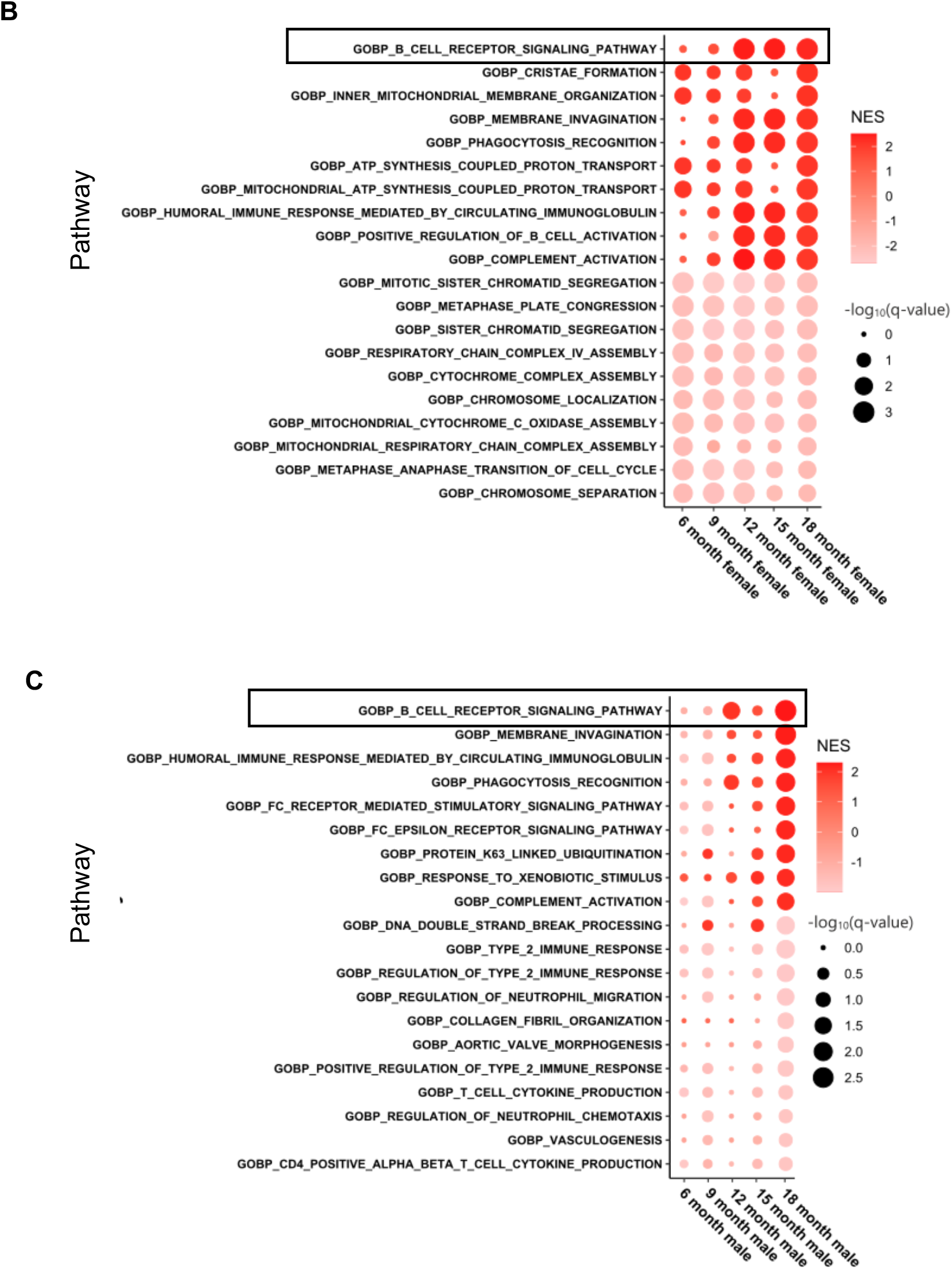
Urinary bladder of healthy C57BL/6 mice exhibit age and sex-associated enrichment in immune response pathways. Enrichment of the top 10 significantly enriched and downregulated GO biological processes pathways in **A)** females compared to age-matched males, **B)** aged females compared to 3-month-old females and, **C)** aged males compared to 3-month-old males. Total RNA from healthy male and female urinary bladders from different age groups was subjected to bulk-RNA sequencing. Differential expression was assessed using DESeq2 in R (1.4.0). GSEA pre-ranked analysis of whole transcriptomic Log2foldchange ranked lists from DESeq2 was performed. Plots were generated using ggplot2 (3.3.5), with dot size corresponding to -log10(q-value, FDR), and the color associated with normalized enrichment score (NES). Black boxes highlight the B cell receptor signaling pathway.

**Figure 2.**
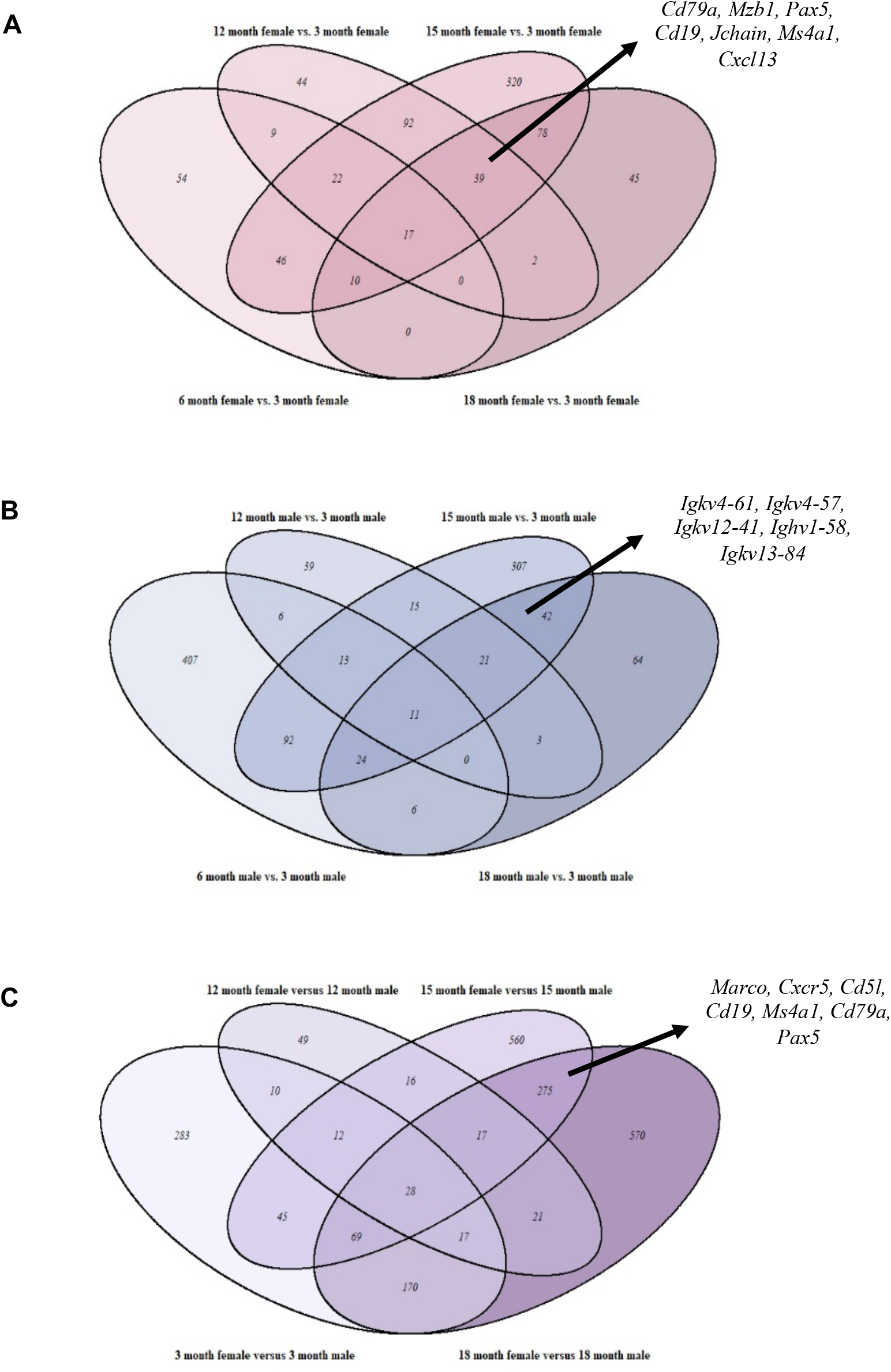
Sexual dimorphism in transcriptomic profiles of aging murine urinary bladders. Venn diagrams showing age and sex associated overlaps and intersecting transcriptomic profiles of significantly differentially upregulated genes in aged mice compared to 3-month-old mice within **A)** females **B)** males, and between **C)** age-matched males and females. Examples of genes associated with B cells are indicated by arrows. Total RNA from healthy male and female urinary bladders was subjected to bulk-RNA sequencing. Differential expression was assessed using DESeq2 program in R (1.4.0) with a *P-* adj < 0.05 (FDR) and a log2fold change cut off of 0.58. Venn diagrams were generated from resulting gene lists using the VennDiagram (1.6.2) package in R.

As illustrated by the Venn diagram of gene lists significantly upregulated in 6-, 12-, 15- and 18-months old females compared to the 3 month old females (log2fold change cutoff of 0.58, adjusted P value (*P*-adj) <0.05), B-cell associated genes (*Cxcr5, Cxcl13, Cd19, Cd79a, Pax5, Mzb1, Ms4a1;* Supplementary Table 2 and *4*) were commonly differentially expressed in aged females (upregulated in 12-, 15-, 18-month females; Figure 2). Similar analysis within males (Supplementary Table 3) and between males and females (Supplementary Table 4), revealed a comparable age associated relationship, but between 15- and 18-month age groups only. Notably, these overlapping gene lists included a large number of immune response associated genes in aged female mice compared to young mice or age-matched male mice. In addition, Immgen MyGeneset application based immune cell prediction further showed increased shifts in various B cell subsets associated genes initiating at 12 months in females, whereas in males, similar increases were observed at 18 months of age. Between sex comparisons revealed substantial upregulation in B-cell populations in females starting at 15 months (Figure 3A).

**Figure 3.**
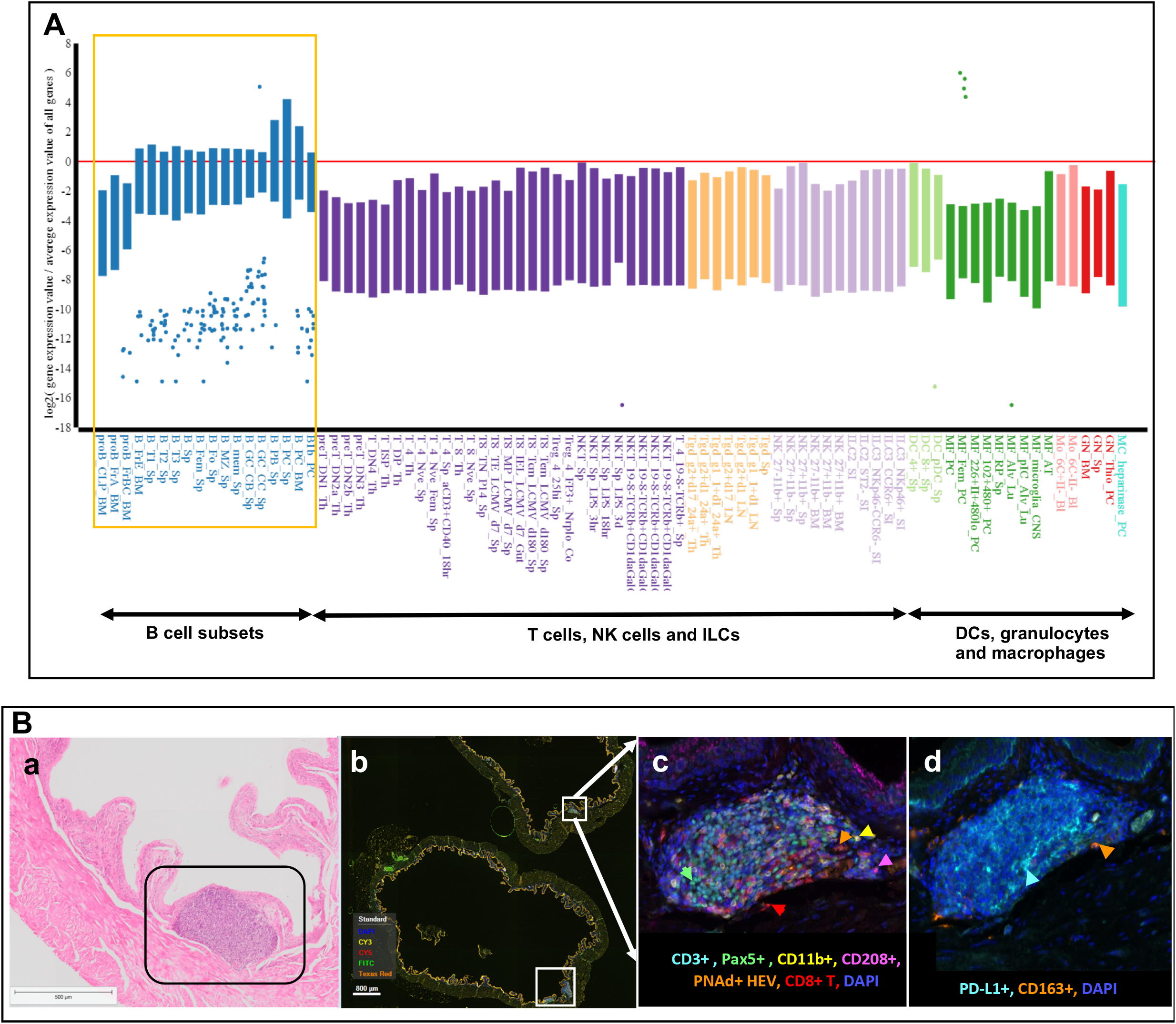
Age associated increase in B cell alterations in female mouse bladder. **A)** Healthy male and female mouse bladders (n=3/4) of ages, 3, 6, 9, 12, 15 and 18 months were subjected to bulk RNA-Seq. Differential expression was assessed using DESeq2 in R (1.4.0) with a *P-*adj < 0.05 (FDR). Analysis of differential expression gene lists was performed using the MyGeneset application at https://www.immgen.org/, to determine immune cell subset type enrichment. Plot represents comparison between top 200 differentially expressed genes between 15-month-old female and age matched males. **B)** Hematoxylin and eosin stained section showing lymphoid aggregate in an 18-month-old female mouse bladder wall **(a)**. Whole bladder image captured using Phenochart Software image viewer 1.0.9 (Akoya Biosciences) from multiplex immunofluorescence stained 18-month-old female bladder section showing tertiary lymphoid structures **(b;** white boxes). Immune cells within TLSs (F4/80+ macrophages, CD163+ M2 macrophages, Pax5+ B cells, CD3+ T cells, CD8+ cytotoxic T cells and PD-L1 immune checkpoint **(c** and **d)**. Representative of n=3-5 in each sex.

### Tertiary lymphoid structure formation occurs in the healthy bladder mucosa with biological aging

Histopathological evaluation followed by multiplex IF showed higher numbers of lymphoid aggregates/TLSs within the lamina propria of aged females (Figure 3B; panel a). In both male and female mice, these aggregates first appeared at 9 months of age and with a larger diameter in female bladders. Multiplex IF staining revealed the presence of B, T, cytotoxic T, CD11b myeloid cells, and high-endothelial venule (HEV) confirming a TLS (Figure 3B; panel c). Only the aged female groups appeared to have organized TLS structures within the lamina propria. Given the regulation of PD-L1 immune checkpoint by X-chromosome associated miRNAs, we also measured its expression in this study. Higher expression of PD-L1 was observed within the TLSs along with higher infiltration of CD163+ cells in 18-month-old mice from both sexes (Figure 3B; panel d). These findings are suggestive of sex differences in mechanisms underlying aging associated increased magnitude of inflammatory changes within the urinary bladder.

### BBN carcinogen treatment leads to distinct age and sex associated immunologic alterations in the bladder microenvironment

Based on the findings from healthy murine bladder transcriptomic profiling, we next sought to define the age group that more closely reflects the systemic and local immunologic functional states at the time of local treatment initiation in NMIBC. Female and male mice belonging to young and older age groups were exposed to BBN carcinogen for 12 weeks. Histopathological evaluation of bladders collected at various time points revealed the presence of reactive atypia at 4 weeks and the presence of urothelial dysplasia at 7 weeks post BBN in the mice from younger age group. Presence of dysplasia with focal carcinoma *in situ* was present at 12 weeks post BBN in 100% of mice in this age group. In the 12 months old male mice, early stages of invasion to lamina propria was present in 3 out of 5 mice at 12 weeks post BBN administration (Figure 4F). In age matched female mice, invasion to lamina propria was observed in one out of 5 mice and remaining 4 exhibited dysplasia with widespread carcinoma *in situ* at this time point (Figure 4J). Interestingly, in the 15-month-old groups, invasion to lamina propria and formation of TLSs was evident at 7 weeks post BBN in both sexes. Intensity of hyperplasia was higher in both sexes in this age group (Suppl. Fig. S4A & S4B). However, the abundance of plasma cells in the lamina propria of the female mice from the older age groups was more pronounced in comparison to young females and males from both age groups in the study (Supplementary Figures 4C and 4D). Another important finding was the higher magnitude of overall immune cell infiltration in the lamina propria, accompanied with edematous changes, in female mice in both young and old age groups (Figure 4A-D). Thickening of the bladder wall due to these changes was also observed via ultrasound imaging (data not shown). Myeloid cells were the most abundant across all age groups and showed presence of both F480+ and CD163+ M2 like macrophages (Figure 4G and 4K). Increased expression of PD-L1 on urothelium and endothelial lining of vessel walls was observed in the lamina propria of bladders at 12 weeks post BBN (Figure 4G). Most importantly and in line with local immune responses at mucosal surfaces, TLS formation as a result of carcinogen exposure was present in the lamina propria of all BBN carcinogen treated mice in all age groups (Figure 4C and 4D; Supplementary Figure 4C and 4D). Overall, these findings indicate an influence of age on tumor progression in both sexes.

**Figure 4.**
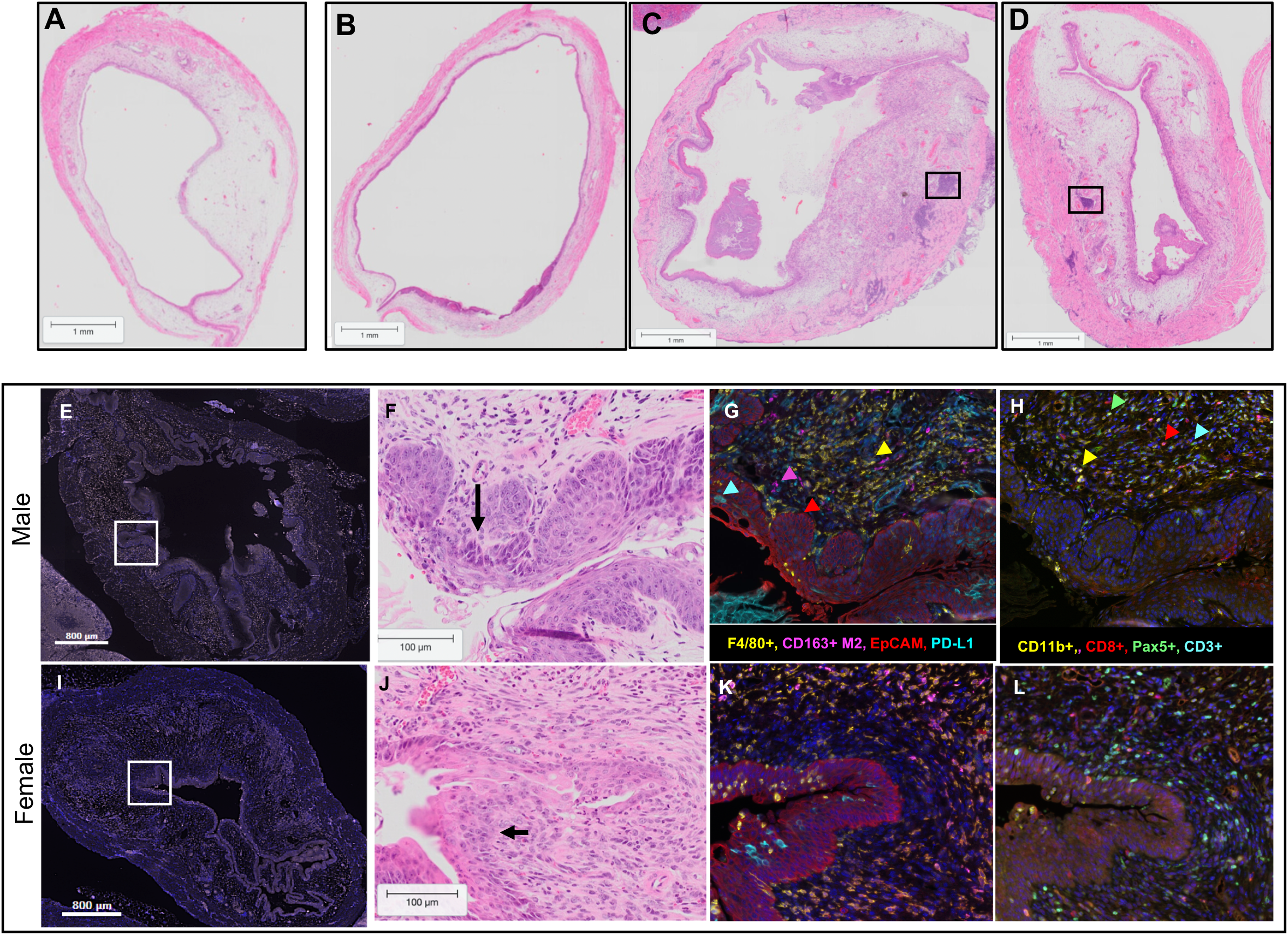
Bladder immune microenvironment following BBN carcinogen exposure. Hematoxylin and eosin (H&E) stained section from 5-month-old female (**A**) and age matched male (**B**) bladder at 12 weeks post BBN exposure. Tertiary lymphoid structure (black boxes) in the lamina propria of 15-month-old female (**C**) and age matched male (**D**) bladders at 7 weeks post BBN. Whole bladder image captured using Phenochart Software image viewer 1.0.9 (Akoya Biosciences) showing low magnification image of multiplex immunofluorescence stained whole bladders from 12-month-old male (**E**) and female (**I**) at 12 weeks post BBN exposure. White boxes indicate representative regions (3-5 per section) selected for high magnification images. Corresponding H&E stained section of a male(**F**) and female (**J**) bladder showing high-grade dysplasia-carcinoma in situ and early invasion. Multiplex immunofluorescence stained image showing F4/80+ macrophages, CD163+ M2 macrophages, PD-L1 immune checkpoint, EpCAM+ epithelial cells (**G** and **K**) and CD11b+ myeloid cells, Pax5+ B cells, CD8+ cytotoxic T cells and CD3+ T cells (**H** and **L**). Representative of n=3-5 in each sex.

## Discussion

Biological aging related sex-disparity in bladder-associated pathologies such as bladder cancer and physiologically healthy bladder mucosal tissue has recently drawn significant attention. Despite discrepancies in the literature and a consensus on age and sex associated hormonal, genetic and immunological deviations, the current pre-clinical models of bladder cancer have yet to fully integrate such factors in experimental design for development of immunotherapeutic agents.

It is known that females usually mount stronger adaptive immune responses to infectious and vaccine associated immune challenges, producing higher antibody titers and more robust innate immune responses. With the similarities in mucosal immune physiology between humans and mice, and the recently reported age-associated, antigenic stimuli-independent inflammation in female murine bladders,^16^ we conducted a comprehensive evaluation of the sex and age-associated bladder transcriptomic and spatial immune profiling in female and male mice. Not surprisingly, an unsupervised feature selection applied to detect differences in transcriptomic profiles of bladders from all age groups led to distinct clustering of groups by age and sex. While this confirmed the unique bladder transcriptome profiles of the groups under study, many of the top upregulated genes revealed shifts in immune-associated genes with increasing age, suggesting that fundamental changes in the aged murine bladder tissue microenvironment were immune-related. These findings are in concordance with the previous report by Ligon et al.,^16^ where such changes were reported in female mice.

Significant enrichment in pathways associated with B cell function was observed in the bladder transcriptome profiles, starting at 9 months of age in both sexes. Evaluation of the top 25% significantly differentially expressed genes in the oldest group revealed increased expression of a substantial number of transcripts associated with a B cell phenotype such as *Cd19, Cd79a, Pax5, Mzb1, Ms4a1 and others.* The age-associated increased B cell density was independently revealed by multiplex spatial immune profiling of the healthy bladder, prominently in aged female mice, that also exhibited an increased prevalence of TLSs. Following birth, such mucosa associated lymphoid tissues (TLSs), consisting of T cells, B cells, macrophages and follicular dendritic cells, develop across various mucosal sites (e.g. respiratory tract, gut, urogenital tract). The neogenesis of TLSs occurs as a result of exposure to normal commensal or pathogenic microbes, IFN-1 activating vaccines that are administered via mucosal routes, or chronic local inflammation. Such induced TLSs generally provide heterologous humoral immune protection at mucosal sites.^25^ As widely reported, TLSs in the bladder also form during the course of normal aging due to increased systemic levels of TNF-α^16^, chronic inflammatory conditions of the urogenital tract such as those observed in interstitial cystitis associated Hunner lesions and bladder cancer.^26–29^

As observed through analysis using the Immgen MyGeneset application, the increased levels of transcripts associated with various B cell subsets are suggestive of multiple populations dispersed within the healthy bladder mucosa that may or may not be intrinsic to TLSs. Interestingly, the immunoglobulin light and heavy chain variable region genes constituted the majority of the top 50 genes distinguishing the profiles of aged female mice from the younger groups. Although such differences in these B cell associated transcripts showed an increasing pattern of abundance with advancing age in both sexes, these observations were found to be more pronounced in bladders from 18 months old female mice compared to all other groups.

In both humans and mice, B cells are the only immune cells that exhibit age-related sexual dimorphism.^12,30^ With advancing immunologic and biologic aging, peripheral pools of B cells show significant decline with a simultaneous increase in mucosal surfaces. In the current study, the *Cxcr5* gene was also significantly overexpressed in the bladders of old females with concurrent higher expression of the gene *Cxcl13* encoding the B-cell recruiting chemokine CXCL13. Higher transcripts levels of *J chain* also indicate a higher amount of secreted IgM and IgA antibodies in the bladder mucosa. Indeed, identification of the functional properties of these cells in the context of carcinogenesis and locally administered BCG immunotherapy in NMIBC is needed to augment the identification of newer therapeutic targets.

Another interesting and novel observation in our study was the increased expression of *Pd-l1* immune checkpoint gene in bladder transcriptome profiles of aged healthy female mice, which was further also observed at the protein level within the TLSs, reflective of exhausted states. Increased expression of other immune checkpoint genes such as *Lag3* and *Btla,* was also observed in the bladders from healthy older females compared to their male counterparts. A bulk sequencing approach does not allow the identification of specific cell types that exhibit such exhausted behavior and thus characterization of cell type specific lack of function via single cell sequencing would allow for a comprehensive understanding of such alterations and development of precise targeting approaches.

In a recent report, Degoricija et al., showed the immunologic alterations and higher expression of immune checkpoint molecules in tumors from young male mice at 20 weeks following exposure to BBN carcinogen.^31^ Although this is one of the few reports that have characterized some local inflammatory changes accompanying carcinogens, this study was conducted in young male mice. Based on our observation on increased PD-L1 protein expression in the urothelial and endothelial cells, it can be speculated that this may result from BBN induced DNA damage and activation of cellular IFN pathways. Such increased PD-L1 on endothelial cells is known to cause inhibition of T cell activation.^32^ Future mechanistic studies targeting this checkpoint are thus warranted to define its dynamic role in disease progression and potential immune exclusion.

Findings from our study showing higher density of plasma cells in the lamina propria of old mice, irrespective of sex, aligns with the increased recruitment of B cells to the bladder mucosa with advancing age that we observed in our findings from healthy bladder molecular profiling. It is plausible that chronic BBN exposure further amplifies the recruitment and differentiation of B cells to plasma cells. This is supported by the observation that increase in immune infiltrates and lymphoid aggregate formation was observed as early as week 4 post BBN initiation in majority of mice in all age groups.

The biological and immunological aging related similarities in humans and mice, emphasize the inclusion of aged mice in pre-clinical studies in bladder cancer. This is more important especially in studies investigating locally delivered BCG immunotherapy in NMIBC to establish the role of resident and treatment induced or recruited immune cells that may exhibit age and sex dependent function in treatment response. Indeed, tumor associated B cells and bladder tumor-associated TLSs are gaining increased attention because of their association with response to BCG (*Unpublished findings*) and positive outcomes following immune checkpoint blockade.^28,33^ Future in-depth investigations into the interactions between these pre-existing TLSs and locally delivered immunotherapy are necessary. Finally, of the other immune cell types such as cytotoxic T cells and myeloid cells, we observed a high density of both F4/80+ and CD163+ M2-like macrophages in the bladder immune microenvironment of BBN treated mice.

The majority of studies in bladder cancer are conducted in 6-8-week-old mice. The influence of hormones on sex differences in time to tumor induction in the BBN model was established in a seminal study over 4 decades ago.^34^ However, the impact of immunologic aging that accompanies hormonal alterations remains to be fully understood. Moreover, mice do not undergo menopause and significant decline in estrogen levels as seen in humans,^35^ and exhibit reproductive senescence associated decline in ovarian function. It is thus challenging to model the cross-talks between hormones and immune alterations precisely. Moreover, the role of androgen and androgen receptors in mediating BBN carcinogen associated sex differences in mice is well established.^36,37^

Overall, our novel findings on age and sex related immune landscapes in physiologically healthy bladder and the potential influence on the immune microenvironment of the BBN carcinogen induced model of bladder cancer suggest that these factors are critical in pre-clinical modeling of the disease. Use of older mice in development of immune potentiating therapies in bladder cancer will facilitate precise translation to humans. Further investigations in the context of immunotherapy response are justified to understand the importance of age and sex in mediating treatment response and overall outcomes.

## Author contributions and acknowledgements

MK conceptualized and designed the overall study. AH conducted healthy bladder RNA-sequencing and wrote the manuscript with MK and DL. DL and KT performed bioinformatics data analysis. GS, SC and MK performed BBN treatment experiments. MX performed histopathological evaluation of the BBN carcinogen exposed murine bladders. PY evaluated all multiplex immunofluorescence imaging data from the BBN model. All authors reviewed and edited the manuscript. Katy Milne at BC Cancer’s Molecular and Cellular Immunology Core Histology department at the Trev and Joyce Deeley Research Centre helped with the antibody panel design and multiplex immunofluorescence staining and imaging. Funding for this study was provided by Mary and Mihran Basmajian award for Excellence in Health Research, Queen’s University and Bladder Cancer Canada to MK.

## Supplementary methods

### RNA-sequencing and data analysis

Libraries were prepared with a Ribo-zero rRNA depletion kit (Illumina) followed by sequencing using the illumina NovaSeq 6000 S4 PE100 platform. Quality of Fastq files was assessed using FastQC, and the presence of adapter sequences was observed. Reads were trimmed using Trimmomatic (0.36) using the following reference adapters; Forward: AGATCGGAAGAGCACACGTCTGAACTCCAGTCAC – Reverse: AGATCGGAAGAGCGTCGTGTAGGGAAAGAGTGT. Fastq files were then pseudoaligned against the mouse transcriptome (GRCm38.p6) from Gencode Release M25 using Salmon (1.3.0) in mapping mode with co-dependencies nixpkgs (16.09), openmpi (3.1.2), and gcc (7.3.0) on the Queen’s Centre for Advanced Computing cluster. This transcriptome is in ENSEMBL annotation. Raw and TPM normalized counts from salmon Quant.sf files were then extracted into R (4.1.0) using Tximport to get read coverage tables. TPM counts were visualized using Matlab R2020b for preprocessing. Differential expression of raw counts was performed then performed in R 4.1.0 using DESeq2. Genes were ranked by Log2foldchange for within sex age associated differences and between sex pairwise comparisons for all age groups.

Given that GSEA uses human gene sets, the ranked lists were collapsed and remapped using the Mouse_ENSEMBL_Gene_ID_Human_Orthologs_MSigDB.v7.4.chip before GSEA pre-ranked was run for 1000 permutations, with inclusion criteria for gene sets set from 15 to 200.^21^ Results were visualized through dot plots generated with ggplot2 (3.3.5) in R (1.4.0) representing the top 10 upregulated and downregulated GO Biological processes pathways in aged versus young female and male mice and age-matched females vs. males. Results of all significantly upregulated and downregulated pathways were also visualized using Cytoscape (3.8.2; http://www.cytoscape.org) based on the workflow developed by Reimand et al.^21^ Briefly, Enrichmentmap was run with FDR q-value < 0.05 and combined coefficient >0.375 with combined constant = 0.5 within Cytoscape. Nodes were clustered and labelled using AutoAnnotate run with the default MCL cluster algorithm with similarity coefficient edge weight column and a maximum of 10 annotations before being manually arranged and renamed to represent major biological themes. Given that genes were in ENSEMBL annotation in the TPM and raw data tables, genes from raw and TPM read tables with a known name were extracted and renamed using the org.Mm.eg.db package in R Bioconductor (3.13). DESeq2 was rerun using this raw data for all comparison groups and filtered with false discovery rate (FDR) adjusted *P*-value <0.05 (false discovery rate). Venn diagrams were generated using the VennDiagram packaged (1.6.2) in Rto reveal the overlaps between profiles of significantly overexpressed genes (*P*-adj <0.05, FDR with a log2foldchange cut off of 0.58) that associated with age and sex.

**Supplementary Table 1.** Antibodies used in multiplex immunofluorescence staining.

| **Primary Antibody** | **Staining Round** | **Species** | **Cat#** | **Clone #** | **Ab Dilution (Diluent)** | **Fluorophore** | **Fluorophore dilution** |
| --- | --- | --- | --- | --- | --- | --- | --- |
| CD208 | 1 | Rat | DDX0191P-100 | 1010E1.01 | 1/100 RR | Opal 650 | 1/100 |
| CD3 | 2 | Rabbit | M3074 | SP7 | 1/600 DVG | Opal 520 | 1/300 |
| PNAd | 3 | Rat | 120802 | MECA-79 | 1/200 DVG | Opal 620 | 1/100 |
| Pax5 | 4 | Rabbit | ab109443 | EPR3730 | 1/1000 DVG | Opal 540 | 1/500 |
| CD8 | 5 | Rabbit | 98941S | D4W2Z | 1/200 DVG | Opal 690 | 1/150 |
| CD11b | 6 | Rabbit | ab133357 | EPR1344 | 1/7500 DVG | Opal 570 | 1/800 |
| CD163 | 1 | Rabbit | ab182422 | EPR19518 | 1/3000 RR | OPAL570 | 1/200 |
| Ly6G | 2 | Rat | 127607 | 1A8 | 1/800 DVG | OPAL620 | 1/300 |
| EpCAM | 3 | Rabbit | ab221552 | EPR20533-63 | 1/600 DVG | OPAL690 | 1/100 |
| PD-L1 | 4 | Rabbit | 64988 | D5V3B | 1/150 DVG | OPAL520 | 1/100 |

