## Supplementary Figures for "The influence of age and sex on the pre-treatment immune microenvironment of a carcinogen induced murine model of bladder cancer"

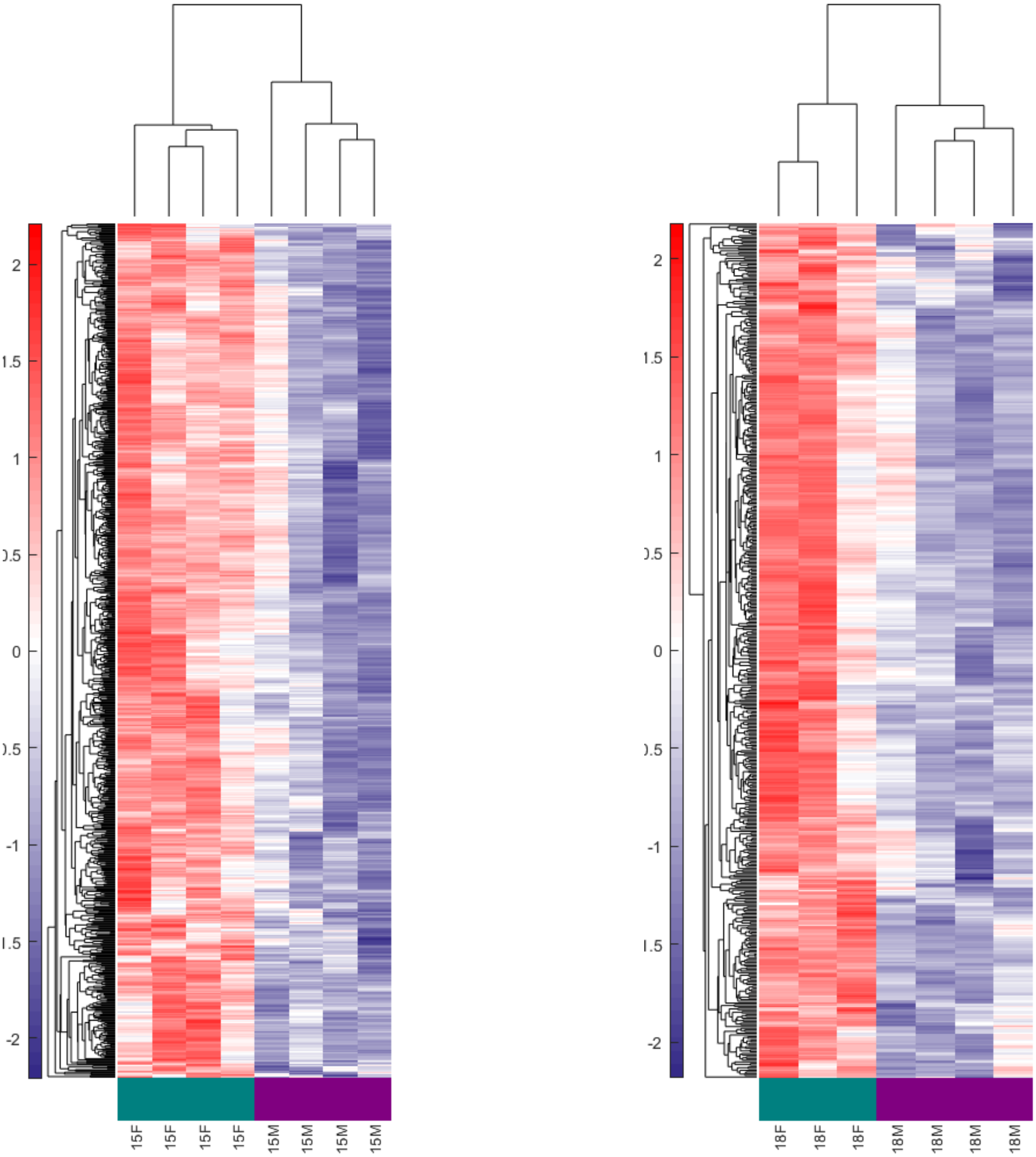


**Supplementary Figure 1. Unsupervised log2 mean-centered heatmap of RNA sequencing data of the top 25% ranked genes between and within sexes** Total RNA from healthy male and female (M=male, F=female) urinary bladders from different age groups was subjected to bulk-RNA sequencing. The top 10% discriminating genes were ranked by an unsupervised feature selection algorithm. Heatmap was generated by setting a threshold of the top 25% ranked genes (significance was determined using MATLAB based feature selection algorithm). Rows represent the relative mean log2 fold change in sample (n=3-4 per age group in each sex).

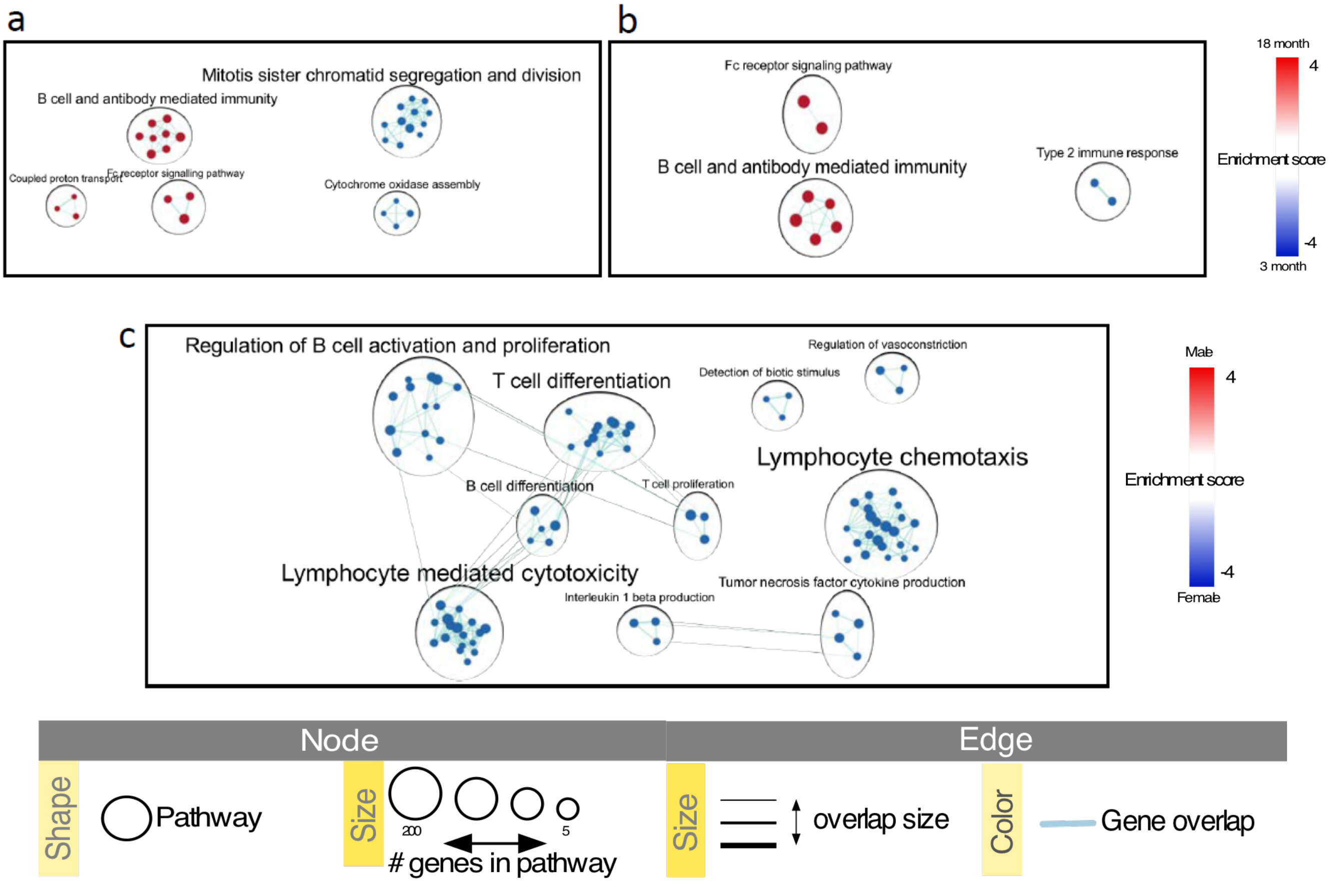


**Supplementary Figure. 2 C57BL/6 mice show increased immune associated pathway enrichment in an age- and sex-dependant manner.** Enrichmentmap was created with FDR *q*-value < 0.05 and combined coefficient >0.375 with combined constant = 0.5 for **a)** 18-month vs 3-month-old females **b)** 18-month vs 3-month old males, or **c)** 18-month-female vs 18-month-old males. Specifically, nodes represent differentially expressed pathways from GO biological processes curated lists and were manually arranged before clustering and labelling using AutoAnnotate within Cytoscape 3.8.2. Autoannotate was run with the default MCL cluster algorithm with a similarity coefficient edge weight column, with a maximum number of 10 clusters. Individual node labels were removed, and clusters were adjusted manually. Clusters were renamed to represent major biological themes. Legend was adapted from Reimand et al.,[21]and edited in Inkscape 1.1.

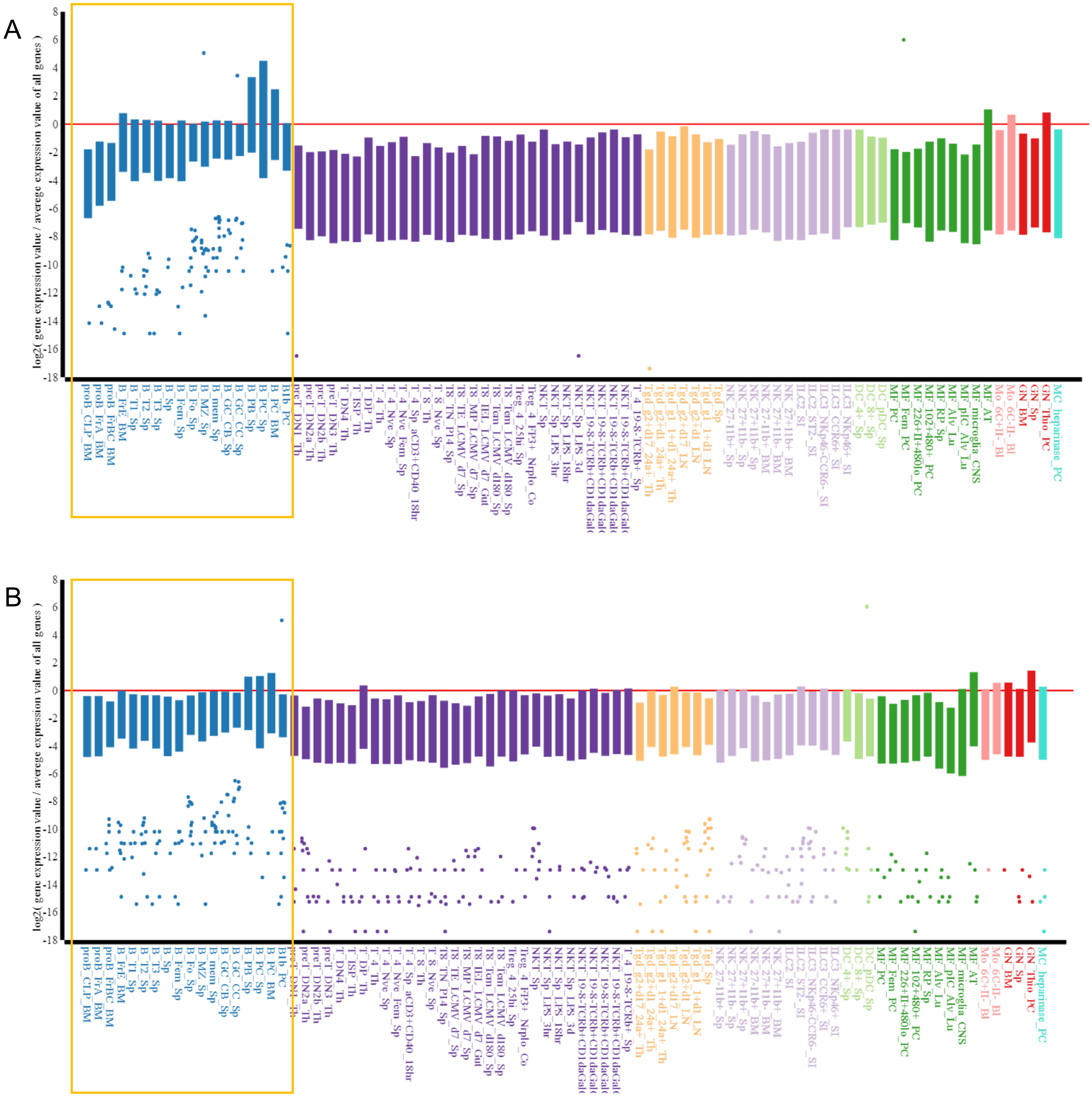


**Supplementary Fig. 3. Age and sex related shifts in B cells in healthy murine urinary bladders.** Healthy male and female mouse bladders (n=3/4) of ages, 3, 6, 9, 12, 15 and 18 months were subjected to bulk RNA-Seq. Differential expression was assessed using DESeq2 in R (1.4.0) with a *P*-adj < 0.05 (FDR). Analysis of differential expression gene lists was performed using the MyGeneset application at https://www.immgen.org/, to determine immune cell subset type enrichment. Plots represent comparison between top 200 differentially expressed genes (FDR *P*-adj<0.05) in 12-month vs 3-month-old female (**A**) and 18-month vs 3-month-old males (**B**).

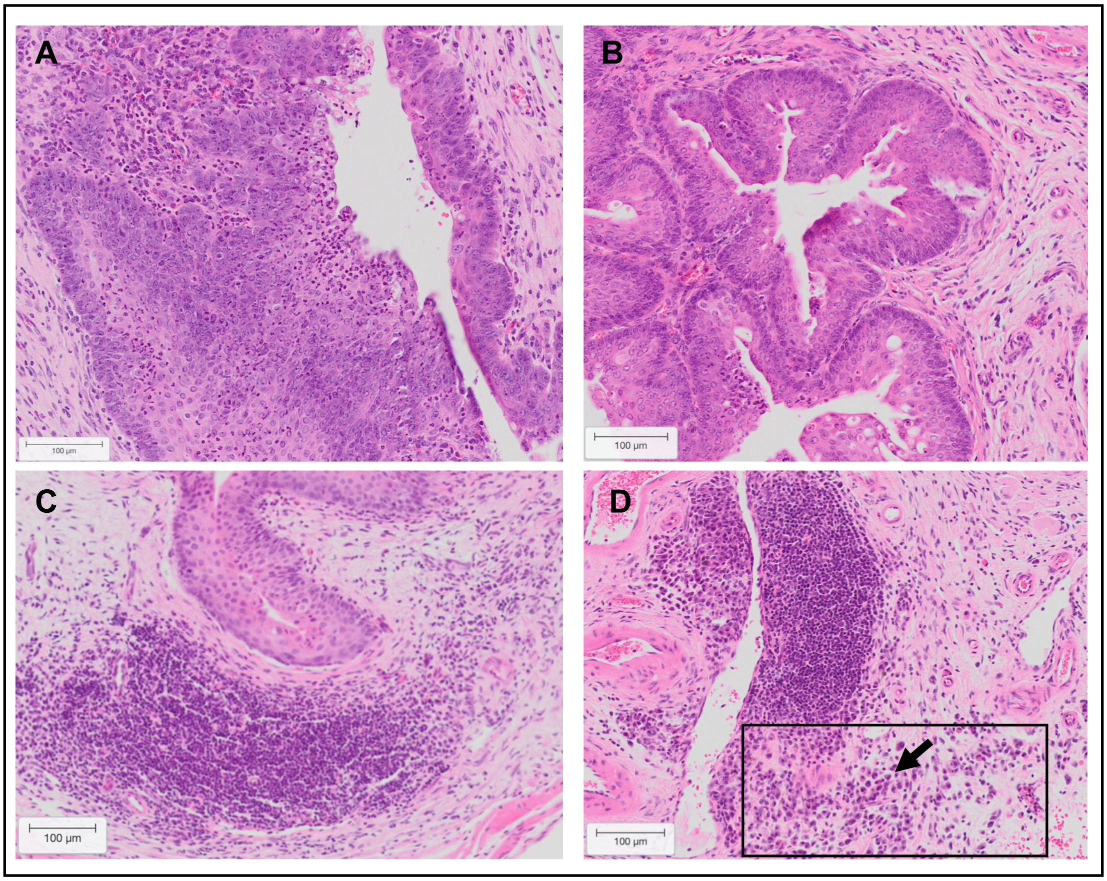


**Supplementary Fig. 4. Bladder microenvironment following exposure to BBN carcinogen.** Hematoxylin and eosin-stained section showing the degree of urothelial hyperplasia and dysplasia at 12 weeks post BBN in a 15-month-old male (A) and female (B) bladder. Tertiary lymphoid structure in the lamina propria of the bladder at week 7 (A) and week 12 (B) post BBN carcinogen exposure in a 12-month-old female. Arrow indicates plasma cells. Representative of n=5 per group

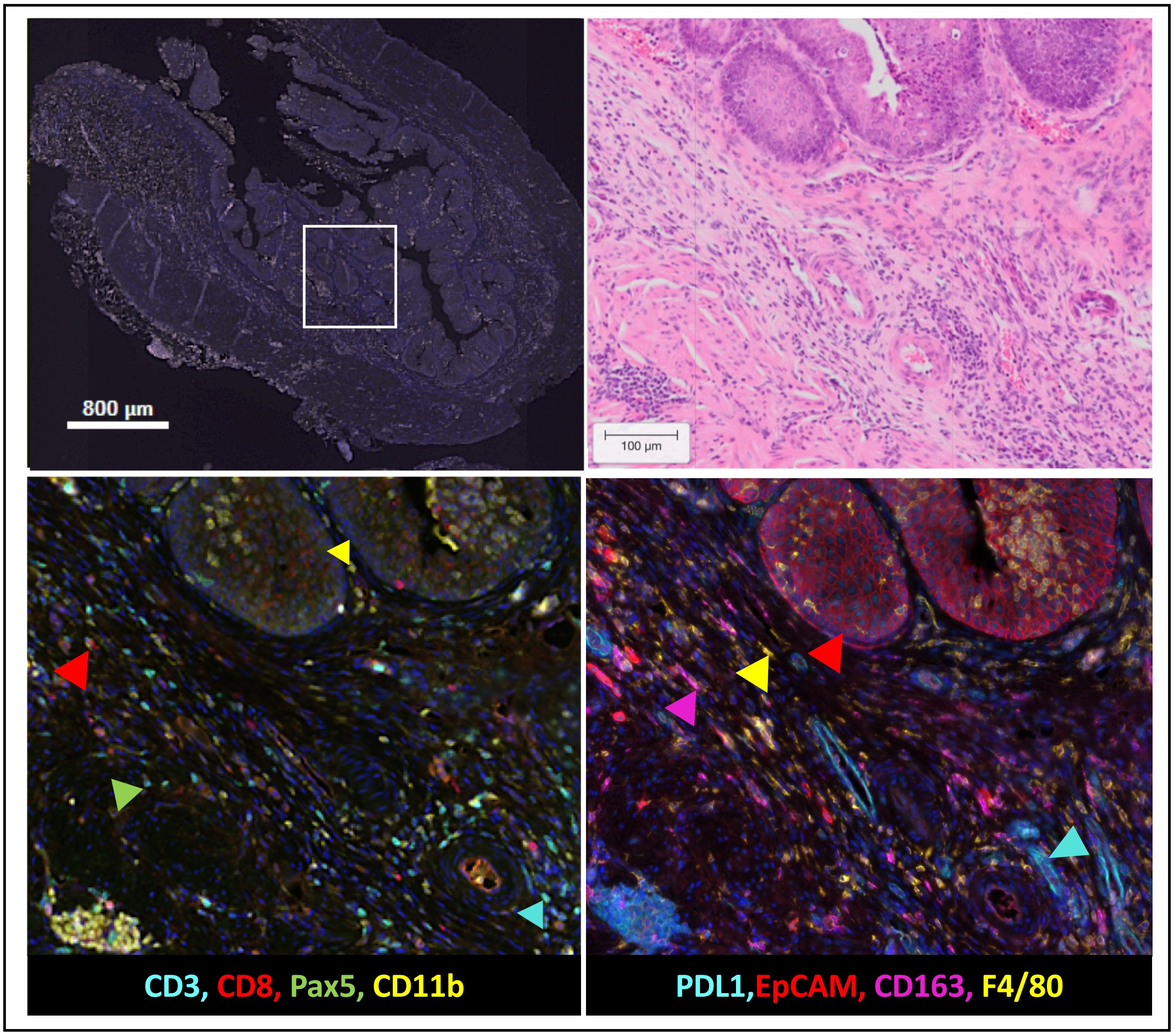


**Supplementary Fig. 5. Immune microenvironment of a 15 month old female bladder following BBN carcinogen exposure for 12 weeks. A)** Whole bladder image captured using Phenochart Software image viewer 1.0.9 (Akoya Biosciences) showing low magnification image of multiplex immunofluorescence-stained whole bladders from 15-month-old female mouse at 12 weeks post BBN exposure. White box in A indicates representative region selected for high magnification images (representative of 3-5 regions selected per section). **B)** Corresponding Hematoxylin and Eosin (H&E) stained section from 15-month-old female bladder at 12 weeks post BBN exposure. Multiplex immunofluorescence stained image showing **C)** CD11b+ myeloid cells, Pax5+ B cells, CD8+ cytotoxic T cells and CD3+ T cells and **D)** F4/80+ macrophages, CD163+ M2 macrophages, PD-L1 immune checkpoint, EpCAM+ epithelial cells. Representative of n=5.

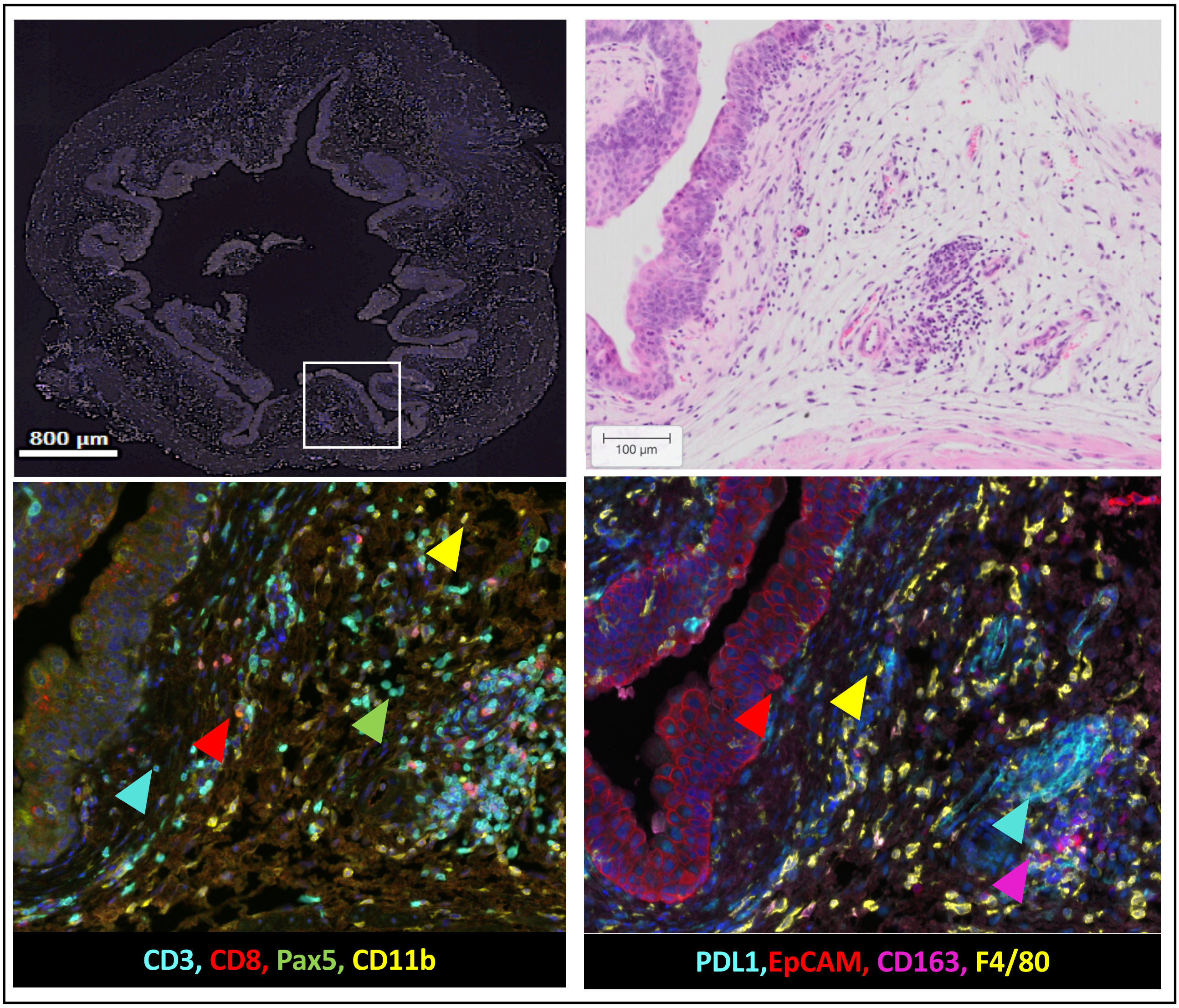


**Supplementary Fig. 6. Immune microenvironment of a 15 month old male bladder following BBN carcinogen exposure for 12 weeks. A)** Whole bladder image captured using Phenochart Software image viewer 1.0.9 (Akoya Biosciences) showing low magnification image of multiplex immunofluorescence-stained whole bladders from 15-month-old male mouse at 12 weeks post BBN exposure. White box in A indicates representative region selected for high magnification images (representative of 3-5 regions selected per section). **B)** Corresponding Hematoxylin and Eosin (H&E) stained section from 15-month-old female bladder at 12 weeks post BBN exposure. Multiplex immunofluorescence stained image showing **C)** CD11b+ myeloid cells, Pax5+ B cells, CD8+ cytotoxic T cells and CD3+ T cells and **D)** F4/80+ macrophages, CD163+ M2 macrophages, PD-L1 immune checkpoint, EpCAM+ epithelial cells. Representative of n=5.
